# Diet composition and sterilization modifies intestinal microbiome diversity and burden of Theiler’s virus infection-induced acute seizures

**DOI:** 10.1101/2023.10.17.562694

**Authors:** Dannielle K. Zierath, Stephanie Davidson, Jonathan Manoukian, H. Steve White, Stacey Meeker, Aaron Ericsson, Melissa Barker-Haliski

**Author notes:** Data Availability Statement: Data from this study will be made freely available by contacting the corresponding author. Ethics Approval Statement: This work was approved by the University of Washington Institutional Animal Care and Use Committee (protocol #4387-02).

## Abstract

**Objective:** Central nervous system infection with Theiler’s murine encephalomyelitis virus (TMEV) in C57BL/6J mice can model acquired epileptogenesis. Diet alters the acute seizure incidence in TMEV-infected mice; yet it is unclear whether intestinal dysbiosis may also impact acute or chronic behavioral comorbidities. This study thus assessed the impact of diet sterilization in a specific pathogen-free vivarium on acute seizure presentation, the composition of the gut microbiome, and chronic behavioral comorbidities of epilepsy.

**Methods:** Baseline fecal samples were collected from male C57BL/6J mice (4-5 weeks-old; Jackson Labs) upon arrival. Mice were randomized to either autoclaved (AC) or irradiated (IR) diet (Prolab RMH 3000 – UU diets) or IR (Picolab 5053 – UW IR diet). Mice then underwent intracerebral TMEV or PBS injection three days later. Fecal samples were collected from a subset of mice at infection (Day 0) and Day 7 post-infection. Epilepsy-related working memory deficits and seizure threshold were assessed 6 weeks post-infection. Gut microbiome diversity was determined by 16S rRNA amplicon sequencing of fecal samples.

**Results:** TMEV-infected mice displayed acute handling-induced seizures, regardless of diet: 28/57 UW IR (49.1%), 30/41 UU IR (73.2%), and 47/77 UU AC (61%) mice displayed seizures. The number of observed seizures significantly differed: UW IR mice had 2.2±2.8 seizures (mean±standard deviation), UU IR mice had 3.5±2.9 seizures, and UU AC mice had 4.4±3.8 seizures during the 7-day monitoring period. The composition of the gut microbiome significantly differed in TMEV-infected mice fed the UU AC diet, with most measured differences occurring in Gram-positive bacteria. TMEV-infected mice fed the UU AC diet displayed worsened chronic working memory.

**Significance:** Intestinal dysbiosis evokes stark differences in acute seizure presentation in the TMEV model and vastly influences the trajectory of post-TMEV infection-induced behavioral comorbidities of epilepsy. Our study reveals a novel disease-modifying contribution of intestinal bacterial species after TMEV-induced acute seizures.

## Introduction

Epilepsy is an underrecognized long-term complication of central nervous system (CNS) infection.^1^ Roughly 19,000 individuals are hospitalized with viral encephalitis annually in the United States alone,^2^ with encephalitis events being highly linked to future development of epilepsy.^3^ Higher incidence of epilepsy in lower income countries (>75% of the 65 million people worldwide with epilepsy) may, in part, be attributable to an increased incidence of CNS infections in these regions.^2^ For example, CNS infection accounts for approximately 14.8% of newly diagnosed epilepsy in Ecuador,^4^ whereas in the United States, CNS infection accounts for only 3% of new epilepsy cases.^5^ Infectious agents are the causative factor in 1% of cases of epilepsy in Ethiopia, but up to 47% of cases in Mali, ^6^ reflecting the highly variable nature of epilepsy of infectious etiology. Viral encephalitis-induced seizures and an increased risk for epilepsy have even recently emerged as a potential secondary outcome of COVID-19 infection, particularly in non-hospitalized children under 16, ^7; 8^ shining a spotlight on the understudied role of viral infections on the risk for seizures and epilepsy. Patients with viral encephalitis who present with seizures are at substantially higher risk (∼22-fold greater) of developing spontaneous recurrent seizures (SRS) post-infection.^9; 10^ Encephalitis itself can lead to drug-resistant epilepsy (DRE).^11^ Infection-induced CNS encephalitis is thus a significant and specific risk factor for acquired DRE, which may indirectly increase the global burden of epilepsy.

Central nervous system infection of C57BL/6J mice with the Theiler’s murine encephalomyelitis virus (TMEV) has emerged as a unique translational model of immune system-mediated acute seizures and epilepsy.^12–17^ As a result, this model carries significant potential to identify untapped mechanisms of ictogenesis and epileptogenesis following viral infection of the brain.^12–17^ Central infection with TMEV in male C57BL/6J mice leads to robust infiltration of pro-inflammatory macrophages, which promotes the upregulation of IL-6 expression necessary for the acute infection-induced seizures in this model.^18^ The pro-inflammatory cytokine TNF-alpha has been previously found to contribute to the onset of TMEV-infection evoked acute seizures.^19^ As a result, pharmacological agents that modulate immune system activation are particularly impactful on acute seizure burden and chronic disease course. We have previously demonstrated that the repurposed anti-inflammatory agent minocycline does not affect acute seizure burden but does blunt the onset of spontaneous recurrent seizure (SRS)-associated behavioral comorbidities (e.g., reduced seizure threshold and increased anxiety-like behavior^17^). Minocycline may exert a disease-modifying effect in the TMEV model through indirect modulation of innate immune system activation during the acute infection period (days 0-10 post-infection; DPI)^20^. Agents that directly modulate macrophage activity (i.e., clodronate) have also been found to significantly alter acute seizure severity in this model.^12^ Neuroinflammation contributes to the development of human epilepsy^18; 21–23^. In this regard, the TMEV model is uniquely positioned to interrogate pharmacological and non-pharmacological interventions that can directly or indirectly impact inflammation. As a result, the TMEV model offers an unparalleled naturally occurring acquired epilepsy model capable of uncovering strategies to modify chronic disease burden.

The gastrointestinal (GI) tract is home to a complex and dynamic bidirectional interaction between the host GI epithelium, host immune system, and myriad resident microbiota that can fluctuate based on host dietary intake or pharmacological history.^24^ The intestinal microbiome is increasingly recognized to modulate the response of the host peripheral immune system. For example, patients with epilepsy who adhere to the ketogenic diet can experience significant shifts in the composition and diversity of intestinal microorganisms.^25^ Long-term probiotic supplementation of patients with drug resistant epilepsy (DRE) can reduce seizure frequency and improve quality of life scores,^26^ whereas short-course ciprofloxacin administration in people with DRE significantly reduces seizure frequency. ^27^ Thus, the precise balance and bacterial composition within the host intestine that is needed to achieve seizure control is presently less clear cut. Chronic stress in mice can also alter the intestinal expression of IL-6 by pro-inflammatory macrophages, but not TNF-alpha or IFN-gamma.^28^ Increased IL-6 expression also activates the STAT3 pathway, which is itself epileptogenic in rodent models of acquired epilepsy.^29^ Thus, intestinal dysbiosis may indirectly increase inflammation and/or lead to an aberrant immune response in the setting of a viral infection that could alter epileptogenesis and/or severity of TMEV-induced epilepsy.

A causal relationship between GI dysbiosis and epilepsy has not yet been definitively established. Sterilization of the diet has been previously found to modify the severity of acute seizures in mice infected with TMEV, however, no causal factor was identified to underlie the differences in acute disease severity.^30^ No study has yet defined whether dietary sterilization alters long-term disease progression and/or behavioral comorbidities after TMEV infection. We thus hypothesized that changes in the gut microbiota coincident with TMEV infection of mice could meaningfully change acute and chronic disease presentation and progression in a specific pathogen-free (SPF) animal facility. Our present study demonstrates that acute TMEV infection and diet sterilization is associated with time-dependent changes in the composition and diversity of the intestinal microbiome, which may altogether influence the severity of acute seizure presentation and chronic disease burden in this mouse model of infection-induced epilepsy.

## Methods

### Animal handling and diet assignment

Male, wild type C57BL/6J, mice (4–5 weeks old; Jackson Labs, Bar Harbor, ME) were group-housed throughout the infection and monitoring period (5/cage). Animals were given free access to assigned diet (below) and reverse osmosis purified, acidified water in bottles except during periods of behavioral manipulation, as previously described.^31^ Animals were maintained in standard housing chambers with corncob bedding in a temperature-controlled SPF vivarium on a 14:10 light/dark cycle (lights on: 6h00, lights off: 20h00) as previously detailed.^31^ For all behavioral studies, mice were allowed to acclimate to the housing facility for at least 5 days and to the testing room for at least 1 hour prior to seizure testing or behavioral manipulation. All studies were conducted between the hours of 9h00 and 17h00 during the animals’ light phase. Animals were euthanized by CO_2_ asphyxiation at the conclusion of the study. This study was not designed to assess the impact of sex as a biological variable or the impact of sex on seizure severity, thus only male mice were used. All animal use was approved by the University of Washington Institutional Animal Care and Use Committee (protocol 4387-02), conformed to the ARRIVE Guidelines,^32^ and was conducted in accordance with the United States Public Health Service’s Policy on Humane Care and Use of Laboratory Animals.

### Fecal sampling

Baseline fecal samples were collected upon arrival from a subset of animals in the study (n = 5/experimental group) and prior to housing within the University of Washington (UW) SPF vivarium. All fecal sampling occurred prior to dietary or infection group randomization and thus reflects the baseline intestinal microbiome biodiversity upon arrival from Jackson Labs and prior to housing in the UW SPF vivarium. Fecal samples were collected coincident with sham or TMEV infection (Day 0) and again on Day 7 p.i., for a total of three fecal samples/mouse (Figure 1) during the acute study period. All fecal samples were collected under sterile conditions.

**Figure 1.**
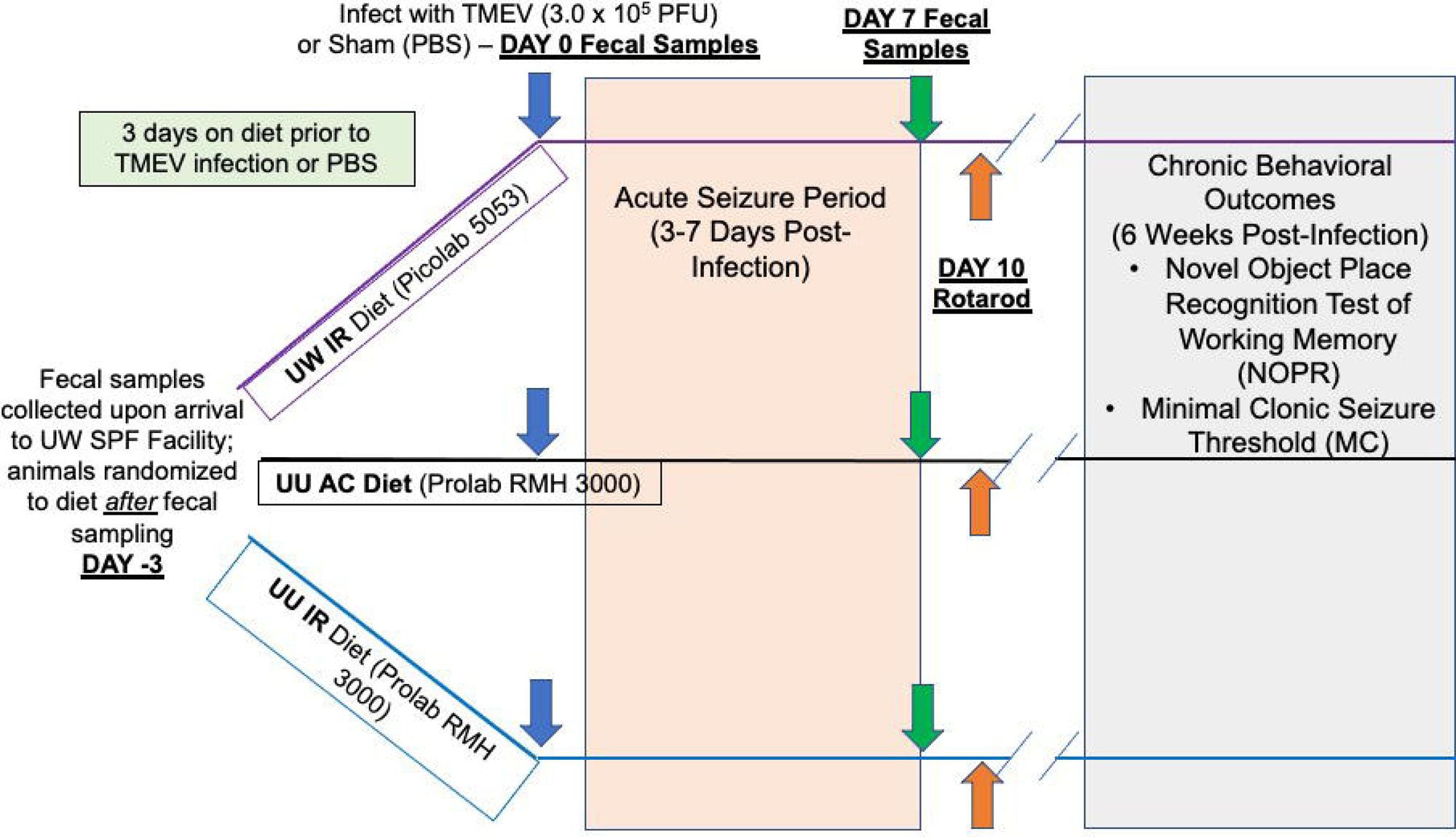
Experimental timeline and study design to assess the effects of dietary sterilization on acute disease course in the TMEV model of infection-induced acute seizures and behavioral comorbidities of epilepsy. Male C57BL/6J mice aged 4-5 weeks old were housed in a specific pathogen-free (SPF) vivarium and provided free access to autoclaved (AC) or irradiated (IR) diet commencing at the time of arrival. Animals remained on specific diet for the duration of the study (to 6 weeks-post infection). Fecal samples were collected at day −3, day 0 (at the time of TMEV infection), and day 7 (end of the acute monitoring period). Mice were assessed twice daily for presence and severity of acute seizures from days 3-7 post-infection; 4-6 weeks later mice were challenged in various behavioral seizure tasks to determine the changes in epilepsy-related behavioral comorbidities.

### Acute TMEV infection and seizure monitoring

Upon arrival, all mice were randomized to receive either an autoclaved (AC) or irradiated (IR) version of the diet used in our prior published studies conducted at the University of Utah^17; 33^ (Prolab RMH 3000), or the house University of Washington diet, irradiated Prolab RMH 5053 (UW IR), with experimental group sizes detailed in Table 1. Animals were maintained on the assigned diet throughout the remainder of the study, including testing of chronic comorbidities 6 weeks after infection. Animals were administered either intracerebral (i.c.) TMEV or PBS (sham) three days later (Day 0). Following TMEV or sham infection, mice were evaluated twice/day for days 3–7 post-infection (p.i.) for the presence and severity of handling-induced behavioral seizures, consistent with our published methods.^17; 33^

**Table 1.** Experimental group sizes for all dietary and TMEV infection groups.

| Infection Group | University of Washington House Diet (Irradiated Picolab 5053) | University of Utah Diet (Irradiated Prolab RMH 3000) | University of Utah Diet (Autoclaved Prolab RMH 3000) |
| --- | --- | --- | --- |
| Sham-infected | 13 | 21 | 26 |
| TMEV-infected ($3.0 \times 10^5$ PFU) | 57 | 28 | 77 |

#### TMEV infection

Mice were free-hand infected intracerebrally (i.c.) with either 20 μL TMEV (titer concentration of 3.0 × 10^5^ plaque-forming units (PFU)) or sterile PBS under isoflurane anesthesia, as previously described.^14; 17; 33–35^. All injection procedures were performed under sterile conditions. Following TMEV injection, the animals were monitored until they had recovered from anesthesia and were ambulatory.

#### Assessment of handling-induced seizures

Mice were evaluated twice/day (minimum 6 hours between sessions), on days 3–7 p.i. for assessment of TMEV-induced acute seizure severity. The presence and severity of handling-induced seizures was scored according to the Racine scale as previously reported.^17; 33; 36; 37^

### Assessment of motor function 10 days p.i. on a fixed speed rotarod

Animals were assessed 10 days p.i. to quantify their recovery of motor function and ability to maintain balance on a rotating rod (6 rpm) for 1 minute over three consecutive trials, as previously described.^17; 38^ The average latency (seconds) to fall off the rotarod was calculated from the three trials, with the group means calculated for each condition.

### Fecal sample 16S rRNA amplicon sequencing

Gut microbiome composition was assessed by 16S rRNA amplicon sequencing of fecal samples from a subset of mice (n = 5 mice/diet) collected at baseline arrival to UW (−3 days p.i.), coincident with TMEV infection (day 0 p.i.), and during active disease (day 7 p.i.).

Fecal samples for 16S rRNA amplicon sequencing were collected at baseline, day 0 p.i., and day 7 p.i. Fecal pellets were collected into individual, sterile tubes, flash-frozen, and stored at −80°C until processing. DNA was isolated from flash-frozen fecal pellets at the University of Missouri Metagenomics Core Center (MUMC) as previously described^39^. Briefly, samples were placed in a 2 mL round-bottom tube containing 800 μL of lysis buffer and a sterile 0.5 cm diameter stainless steel bead and homogenized with a TissueLyser II. Following brief centrifugation to pellet particulates, nucleic acid was extracted using QIAamp PowerFecal Pro DNA extraction kits per the manufacturer’s instructions.

Bacterial 16S rRNA amplicons were generated at the University of Missouri Genomics Technology Core via amplification of the V4 hypervariable region of the 16S rRNA gene using dual-indexed universal primers (U515F/806R) flanked by Illumina standard adapter sequences and the following parameters: 98°C(3:00) + [98°C(0:15)+50°C(0:30)+72°C(0:30)] × 25 cycles +72°C(7:00). Amplicons were then pooled for sequencing using the Illumina MiSeq platform and V2 chemistry with 2 × 250 bp paired-end reads, as previously described.^39^

### Assessment of Chronic Behavioral Comorbidities

Behavioral comorbidities of epilepsy were assessed 6 weeks p.i. for a subset of mice in study (n = 20/TMEV-infection group; n = 10-11 sham) and included: working memory in a novel object/place recognition task and minimal clonic seizure threshold ^40; 41^.

#### Novel object place recognition (NOPR)

This task was conducted based on previously detailed methods.^17; 42^ This task is designed to identify gross changes in working memory in mice with chronic seizures^41^, including in mice previously infected with TMEV ^43^. In a 40 × 40 cm open field chamber each mouse was exposed to a pair of identical “familiar” objects at an approximate distance of 15 cm apart for 15 minutes. The mouse was isolated from the test environment for a 5-minute retention interval while one familiar object was replaced with a novel object in a new location. The mouse was then returned to the testing chamber and exploration behavior was manually recorded for an additional 5 min. The time that the mouse spent with the novel versus the familiar object was determined based on the total time spent with both objects, i.e., the recognition index (RID). A cognitively intact mouse will innately spend more time with a novel object than with the familiar object and thus have a larger RID.^44^ All behavioral evaluations were performed by an experimenter blinded to the treatment condition.

#### Minimal clonic seizure threshold test

Minimal clonic seizure test was assessed after completion of the NOPR task, to assess whether there were long-term reductions in seizure susceptibility between diet and infection groups. Reductions in seizure threshold is a well-established biomarker of epileptogenesis in the TMEV^17; 33^ and other rodent epilepsy models.^45^ Minimal clonic seizures were induced with a 60 Hz, 0.2 second sinusoidal current pulse delivered to the anesthetized (0.5% tetracaine) corneas.^46–49^ These seizures are characterized by rhythmic face and forelimb clonus, rearing and falling, and ventral neck flexion. Mice with these behaviors were considered to have had a seizure. Mice were evaluated repeatedly over the course of 7-10 days, with 48 hours of rest in between electrical stimulation sessions, until group sizes were sufficient to calculate the median convulsant current (CC50) for each treatment condition by the Probit method.^47; 50^

### Statistical Analysis

Daily body weights were analyzed by repeat measures analysis of variance (ANOVA) and post hoc Tukey’s t-test. Latency to first seizure Kaplan-Meier curves was determined with a log-rank (Mantel-Cox) test. Effect of treatment on seizure burden and average number of stage 4/5 seizures were evaluated by the Kruskal-Wallis test. The proportion of severe seizures in each treatment group was determined with a standard score analysis. Bacterial relative abundance data determined via 16S rRNA sequencing were analyzed by two-factor ANOVA. Minimal clonic seizure threshold differences were defined as non-overlapping 95% confidence intervals. Latency to fall off the rotarod, OF measures, and NOPR test results were analyzed using one-way ANOVA and post-hoc Tukey’s t test. All statistical analyses were conducted with Prism v.8.0 or later (GraphPad Software, La Jolla, CA), with p < 0.05 considered statistically significant.

## Results

### Dietary manipulation modifies acute TMEV infection disease presentation

Acute TMEV infection is well-documented to induce acute, handling-induced seizures from roughly day 3-7 p.i.^14; 35; 43^ These seizures in mice are also commonly associated with acute body weight loss and general ill-thrift due to the active viral infection, which can also be exacerbated by acute pharmacological intervention.^17; 33^ Thus, body weight loss was monitored to quantify the extent to which dietary modifications alter metrics of acute symptom presentation post-TMEV infection (Figure 2A and 2B). Total raw body weight was significantly reduced in a time x diet-related manner (Figure 2A; F (30,1231) = 4.26, p < 0.0001) and the percent of body weight change after TMEV infection was also similarly affected by a diet x time interaction (Figure 2B; F(30,1183) = 4.067, p < 0.0001). Thus, dietary sterilization and composition can dramatically influence acute body weight change after a central TMEV infection.

**Figure 2.**
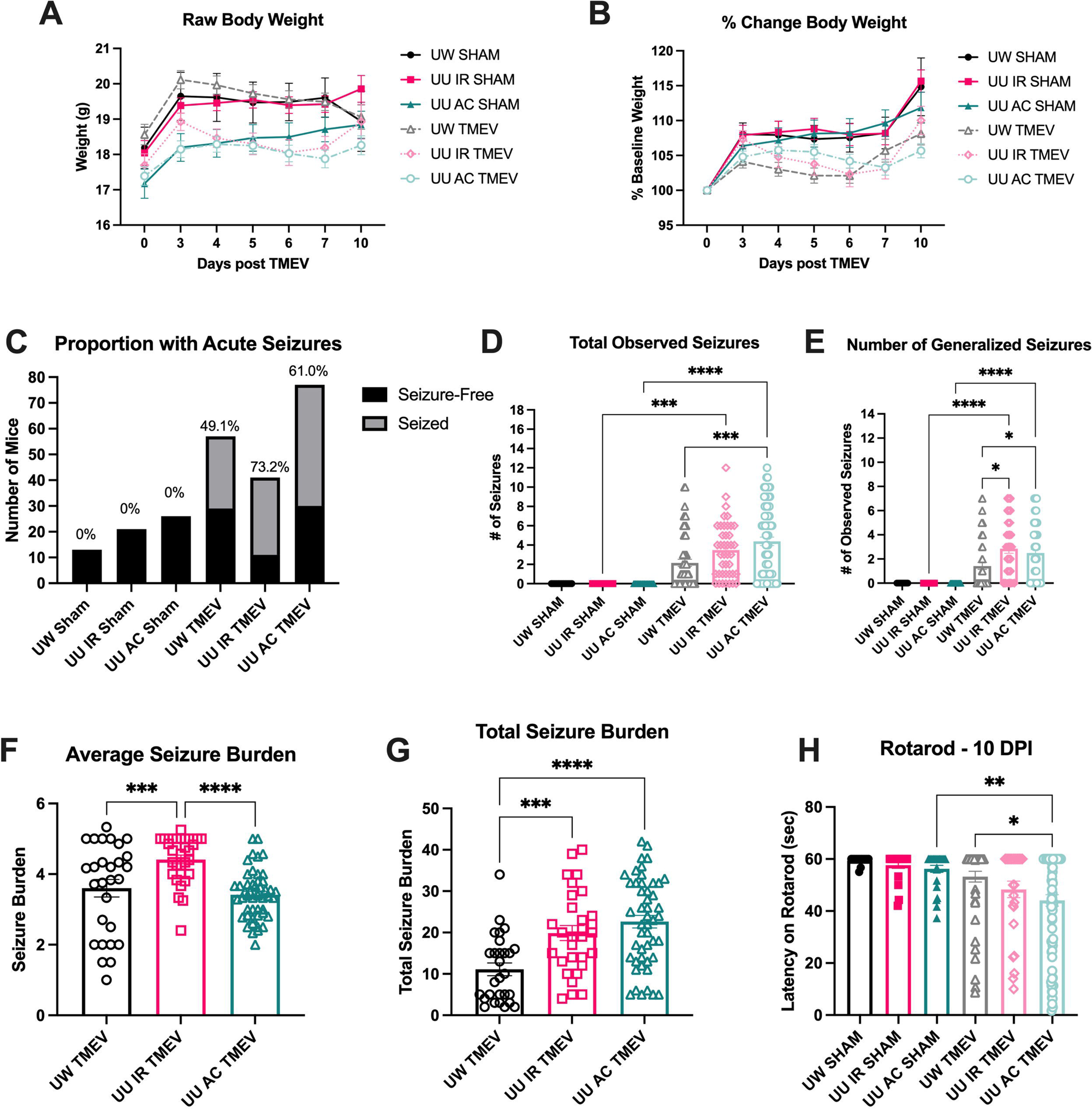
Modulation of diet and diet sterilization significantly altered body weight gain and the acute seizure burden following brain infection with Theiler’s virus (TMEV). A and B) TMEV-induced body weight loss varied with diet (UW versus UU) or diet sterilization procedure (autoclaved – AC; or irradiated – IR). C) Acute seizure incidence is greatest in mice fed an UU IR or UU AC diet. D) The total number of observed seizures during the acute monitoring period is greatest in mice fed UU IR or UU AC diet, but only UU AC diet-fed mice differed from UW diet-fed TMEV-infected animals (p < 0.001). E) The total number of observed generalized seizures (Stage 3 or greater) during the acute monitoring period is greatest in mice fed UU IR or UU AC diet, with both UU IR and UU AC diet-fed mice exhibiting significantly more generalized seizures relative to UW diet-fed TMEV-infected animals (p < 0.05). F) The average seizure burden in mice with any observed seizure was significantly greatest in UU IR diet-fed mice relative to both UW and UU AC-diet fed animals. G) The total seizure burden of mice with any observed seizure was significantly greater in UU IR and UU AC diet-fed mice, but there was no difference in total seizure burden within mice fed the UU diet. H) Recovery from acute infection is only significantly delayed in UU AC-fed mice (p < 0.05).

Libbey and colleagues have previously demonstrated that IR diet cause more mice to develop stage 5 seizures, albeit there was no statistical difference from a non-identical AC diet,^30^ thus we sought to determine whether any treatment condition led to a greater proportion of mice with seizures in mice both administered a matched IR versus AC UU diet as well as whether differences in the diet composition within similar sterilization parameters (UW IR diet versus UU IR diet) could significantly influence the proportion of mice with seizures (Figure 2C). Within the TMEV-infected cohorts, more IR diet-fed mice experienced a Racine stage seizure during the active infection (Figure 2B; X^2^=70.72; p < 0.0001).

TMEV-infection in mice is associated with a high number of acute, handling-induced seizures during the acute infection period (Figure 2D). There was a significant increase in the total number of observed seizures in all TMEV-infected mice (Figure 2D; F (5,229) = 17.09, p < 0.0001). Post-hoc analysis revealed a significant difference in the total number of observed seizures between dietary condition groups, with only UU IR and UU AC diet-fed TMEV-infected mice showing significantly more seizures than diet-matched sham-infected animals (Figure 2D; p < 0.001 and p < 0.0001, respectively. However, only mice fed a UU AC diet demonstrated significantly more observed seizures than the UW IR diet-fed mice, indicating marked differences in seizure presentation within the same institution as a function of diet composition and sterilization. The number of generalized (Racine stage 4/5) seizures was also significantly impacted by diet in TMEV-infected mice (Figure 2E; F(5,229) = 13.35, p < 0.0001), with post-hoc analysis revealing a similar pattern of separation between dietary condition groups after TMEV infection. In this analysis, both UU IR and UU AC diet-fed TMEV-infected mice presented with significantly more handling-induced Racine stage 4/5 seizures than UW IR diet-fed TMEV-infected mice, demonstrating a significant source of potential lab-to-lab variability in the presentation of the TMEV model. Finally, we analyzed both the average seizure burden (Figure 2F) and total seizure burden (Figure 2G) within the TMEV-infected mice that presented with at least one behavioral seizure observed during the acute monitoring sessions. In the TMEV-infected mice with seizures, the average seizure burden was greatest in UU IR diet-fed TMEV-infected mice (Figure 2F) whereas the total average seizure burden was greatest in UU AC diet-fed TMEV-infected mice, suggesting that while there were generally more UU IR TMEV-infected mice with at least one handling-induced seizure, the frequency and severity of seizures was greatest in the UU AC diet-fed animals. Altogether, the acute monitoring period seizure history indicates stark diet-related differences in acute seizure presentation and burden within TMEV-infected mice housed within the same SPF facility.

Finally, performance of TMEV-infected mice with at least one observed behavioral seizure during the 0-8 days p.i. monitoring period were challenged on the fixed-speed rotarod task 10 days p.i. to determine whether there were any differences in recovery from the acute infection at this time point (Figure 2H). Mice infected with TMEV were significantly more impaired on the rotarod than sham-infected mice in the matched diet cohort (F (5, 208) = 5.746, p < 0.0001). Further, post hoc analysis revealed that only UU AC diet-fed TMEV-infected mice experienced significant deficits in motor coordination on this task relative to UU AC diet-fed sham-infected mice, and these animals also exhibited significantly greater impairment on the rotarod task relative to UW IR diet-fed TMEV-infected mice. Altogether, modification of dietary sterilization was associated with substantial changes in the recovery from acute disease between dietary conditions.

### Acute TMEV infection shifts the composition and diversity of the intestinal microbiome by Day 7 post-infection

By Day 7 p.i., at least 49% of all TMEV-infected mice had presented with handling-induced acute seizures (Figure 2C), thus we wanted to ascertain microbial alpha-diversity (Shannon diversity index), as well as relative abundance of specific microbes at this point of active infection relative to that of Day 0 levels (Figure 3). The Shannon diversity index provides a measure within a given sample to indicate both the richness (i.e., number of different taxa found in a sample) and the evenness of distribution of the various taxa, with greater richness and evenness favoring higher alpha-diversity (Figure 3A and 3B). There was no significant difference in Shannon alpha-diversity between diet- and TMEV-infection cohorts at the time of arrival to UW (Figure 3A) yet by Day 7 there was a significant diet × infection interaction on the alpha-diversity of the samples at this time point (Figure 3B; F(2, 44) = 3.243, p = 0.0485). Post hoc analysis further demonstrated that only UU IR diet-fed mice that were infected centrally with TMEV exhibited significant reductions in alpha diversity relative to sham-infected diet-matched cohorts (p = 0.019) whereas UU AC diet-fed mice did not differ by TMEV-infection status, demonstrating that dietary sterilization differences influenced alpha-diversity after central infection with TMEV. These findings altogether demonstrate that dietary modification and sterilization differences within a single facility can dramatically alter intestinal alpha-diversity.

**Figure 3.**
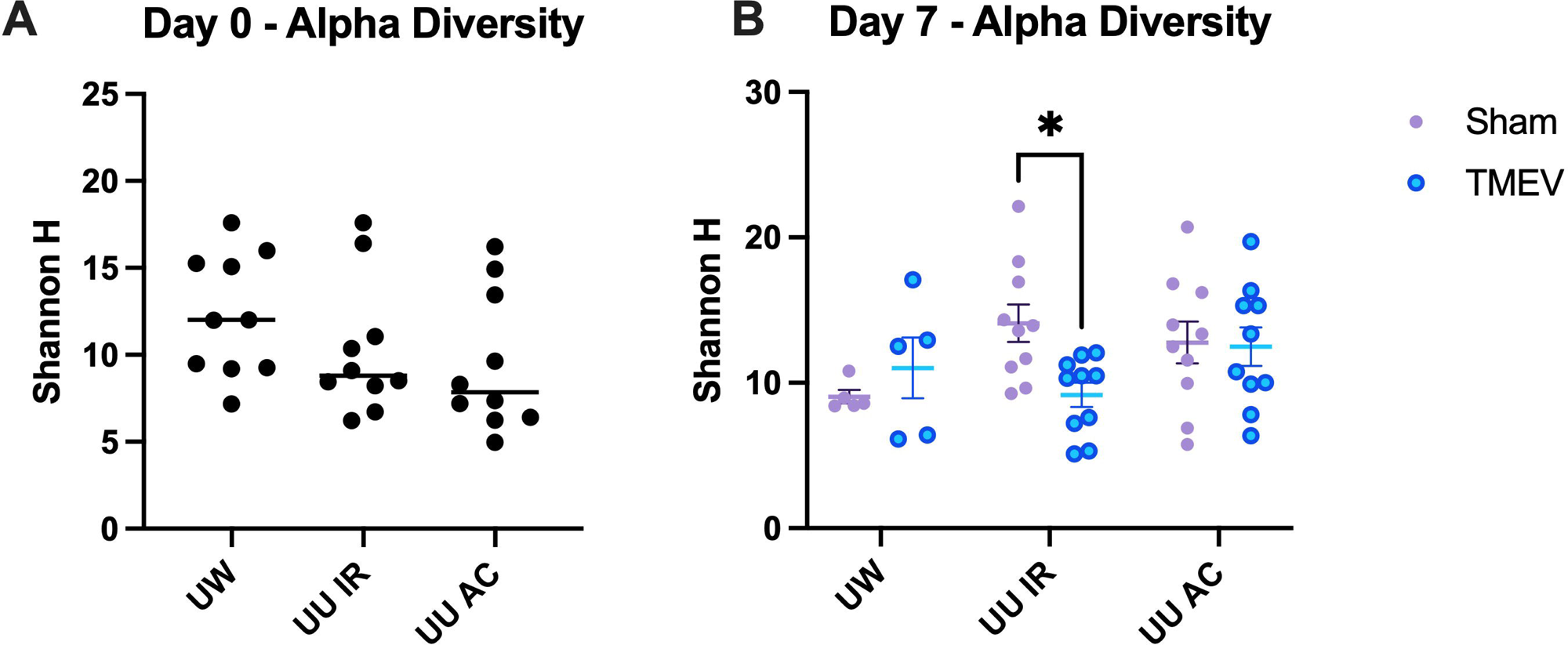
Acute CNS infection with the Daniel’s strain of the Theiler’s murine encephalomyelitis virus (TMEV) model is associated with substantial changes in alpha diversity in the gut microbiome because of dietary composition and sterilization in male C57BL/6J mice. A) There were no significant differences in the relative diversity of the microbiome composition on the day of TMEV inoculation. B) There was a significant diet × infection interaction in respect to alpha-diversity on Day 7 post-infection (F(2,44) = 3.243, p = 0.0485), with post hoc differences evident only in mice that were fed the UU IR diet (* indicates p = 0.0195).

### Diet sterilization vastly alters the composition and diversity of the intestinal microbiome during acute TMEV infection

Intestinal microbial communities are an essential factor underlying many physiological processes including nutrition, inflammation, protection against pathogens, and the function of the nervous system.^51; 52^ In addition to general changes in intestinal biodiversity across time, we also sought to establish whether TMEV infection and dietary modification could specifically and additively alter bacterial species type across the acute TMEV infection period within mice with and without acute handling-induced seizures (Figure 4A-K). The relative abundance of several specific genera of microbes were found to predominate in the fecal matter of mice infected with TMEV or sham (Figure 4). We thus sought to define the extent to which central TMEV infection and modifications of dietary composition could alter the composition and diversity of these specific intestinal microbes in a time-dependent manner (Figure 4A-K). There was a significant time-dependent decrease in the relative abundance of *Akkermansia* (Figure 4A) with time from arrival at our research facilities (main effect of time F(1,726, 41.41) = 65.54, p < 0.0001) but no significant time × diet interaction (F(10,48) = 0.358, p = 0.095). There was a significant time × diet interaction on the relative abundance of *Lactobacillaceae* (Figure 4B) with time (F(10,48) = 4.063, p = 0.0004), with *post-hoc* analysis demonstrating that only the UU AC diet-fed mice infected with TMEV experienced marked shifts in relative abundance of this species relative to UU AC diet-matched sham-infected mice by Day 7 p.i. (p < 0.0001). There was also a time × diet interaction in the relative abundance of *Bacteroidaceae* (Figure 4C; F(10,48) = 2.304, p = 0.0265). *Rikenellaceae* exhibited significant shifts in relative abundance in a time × diet-related manner (Figure 4D; F(10,48) = 3.915, p = 0.0006), but no post hoc differences between groups. There was a main effect of time on the relative abundance of *Muribaculaceae* (Figure 4E; F(1.996,45.91) = 37.79; p < 0.0001), but no post hoc differences. *Bifidobacteriaceae* demonstrated a significant time × diet interaction over the course of the TMEV infection period (Figure 4F; F(10,48) = 2.811, p = 0.008), with post hoc tests revealing marked differences on Day 7 between UU AC diet-fed TMEV-infected mice and diet-matched sham-infected animals (p = 0.015). There was a main effect of housing time on *Erysipelotrichaceae* (Figure 4G; F(1.42,34.07) = 7.147, p = 0.0059), but no post hoc differences. Housing duration also significantly influenced the relative abundance of *Burkholderiaceae* (Figure 4H; F(1.385, 33.23) = 61.82, p < 0.0001), whereas there were no significant effects on *Lachnospiraceae* (Figure 4I). There was a marked time × diet interaction effect on the relative abundance of *Eggerthellaceae* (Figure 4J; F(10,48) = 2.204, p = 0.0336), with post hoc tests indicating that only UU AC diet-fed TMEV-infected mice exhibited significant differences in the abundance of this taxon relative to sham-infected diet-matched mice (p = 0.0015). Lastly, the relative abundance of *Staphylococcaceae* demonstrated a significant time-relative increase (Figure 4K; F(1,24) = 44.05, p < 0.0001) but there were no post hoc differences between groups. Altogether, we presently demonstrate that the relative abundance of multiple intestinal commensals is significantly shifted in a diet- and time-related manner during an acute TMEV infection in C57BL/6J male mice and rehousing in the UW SPF facility.

**Figure 4.**
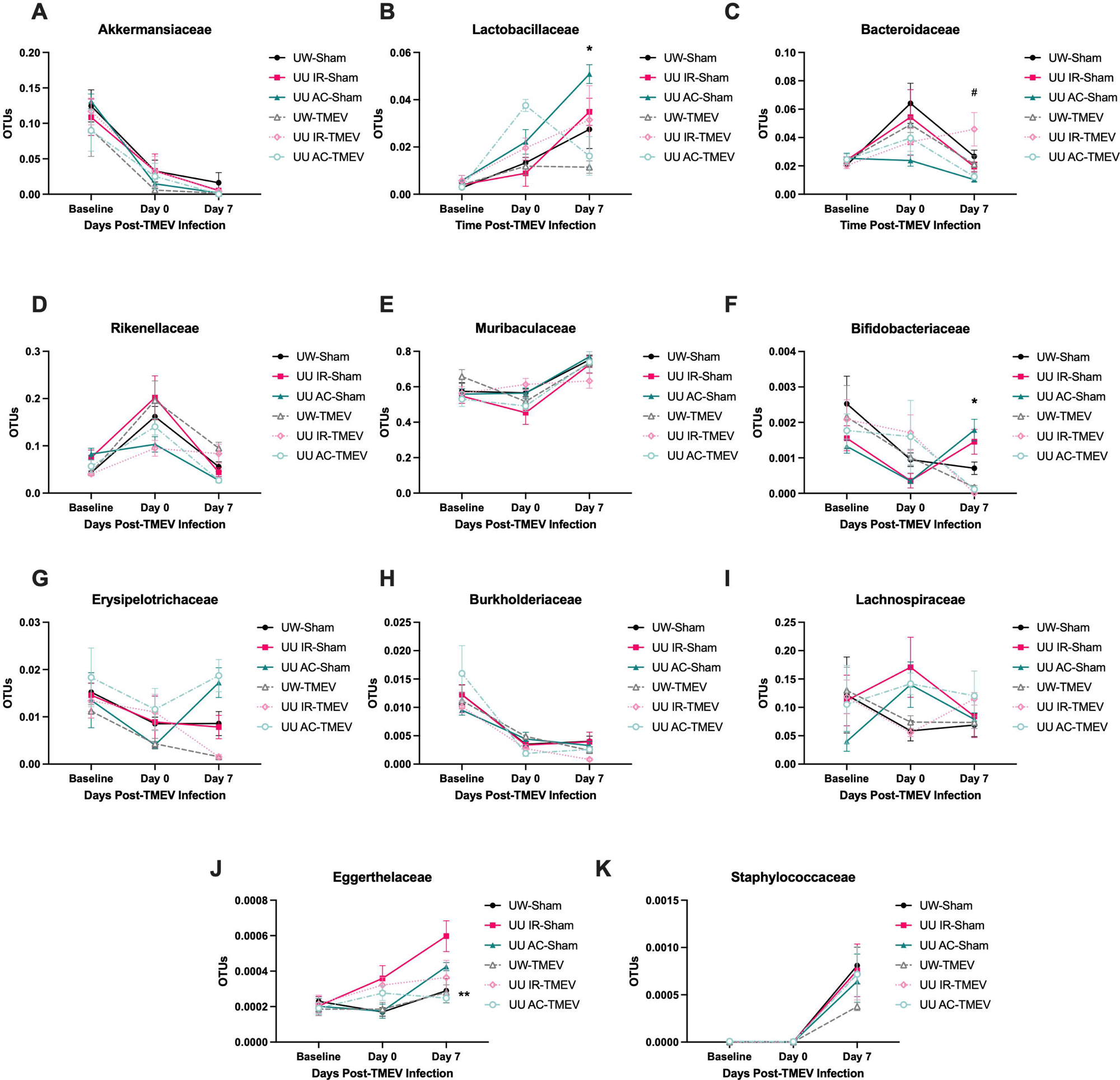
Diet composition and sterilization significantly alters the gut microbial composition during the acute period after CNS infection with the Daniel’s strain of the Theiler’s murine encephalomyelitis virus (TMEV) model in male C57Bl/6J mice. There are marked changes in the phylum-level intestinal microbiome composition and diversity. The most significant changes were observed in Gram-positive bacterial strains, including B) *Lactobacillaceae*, F) *Bifidobacteriaceae*, and J) *Eggerthellaceae.* *indicates significant difference from diet-matched sham-infected mice, p < 0.05. # indicates significant difference between UU AC sham-infected and UW IR sham-infected mice, p = 0.05.

### The composition and diversity of the intestinal microbiome in mice with TMEV infection-induced acute seizures is substantially altered by diet and sterilization

While dietary differences were observed to markedly affect the diversity and composition of the intestinal microbiome during the 10 days from arrival to UW housing facilities, we also sought to assess whether dietary differences in combination with acute seizure history were in any way affecting the diversity and composition of the intestinal microbiome (Figure 5). Thus, the commensal species abundance was assessed within mice that had at least one Racine stage 4/5 seizure versus sham-infected and non-seized mice within each dietary group. The most striking differences between sham-infected and TMEV-infected mice broken down by seizure history were observed only in Gram-positive bacterial species, including *Lactobacillaceae* (Figure 5B; main effect of seizures - F(2,19) = 16.52,p < 0.0001), *Bifidobacteriaceae* (Figure 5F; main effect of seizures - F(2,21) = 22.69, p < 0.0001), and *Eggerthellaceae* (Figure 5J; main effect of seizures – F(2,14) = 3.712, p = 0.050). Gram-positive *Erysipelotrichaceae* exhibited a significant main effect of diet (Figure 5G; F(1,14) = 18.57, p = 0.0007) There were major differences as a function of diet only in the Gram-negative bacterial species, including *Bacteroidaceae* (Figure 5C; F(2,21)=5.174, p=0.0149) and *Rikenellaceae* (Figure 5D; F(2,21) = 10.22, p = 0.0008). Thus, dietary sterilization and composition, coupled with acute seizure history after central TMEV infection leads to marked shifts in bacterial species abundance, with differences in seizure history largely co-occurring with major shifts in gram-positive bacterial species abundance.

**Figure 5.**
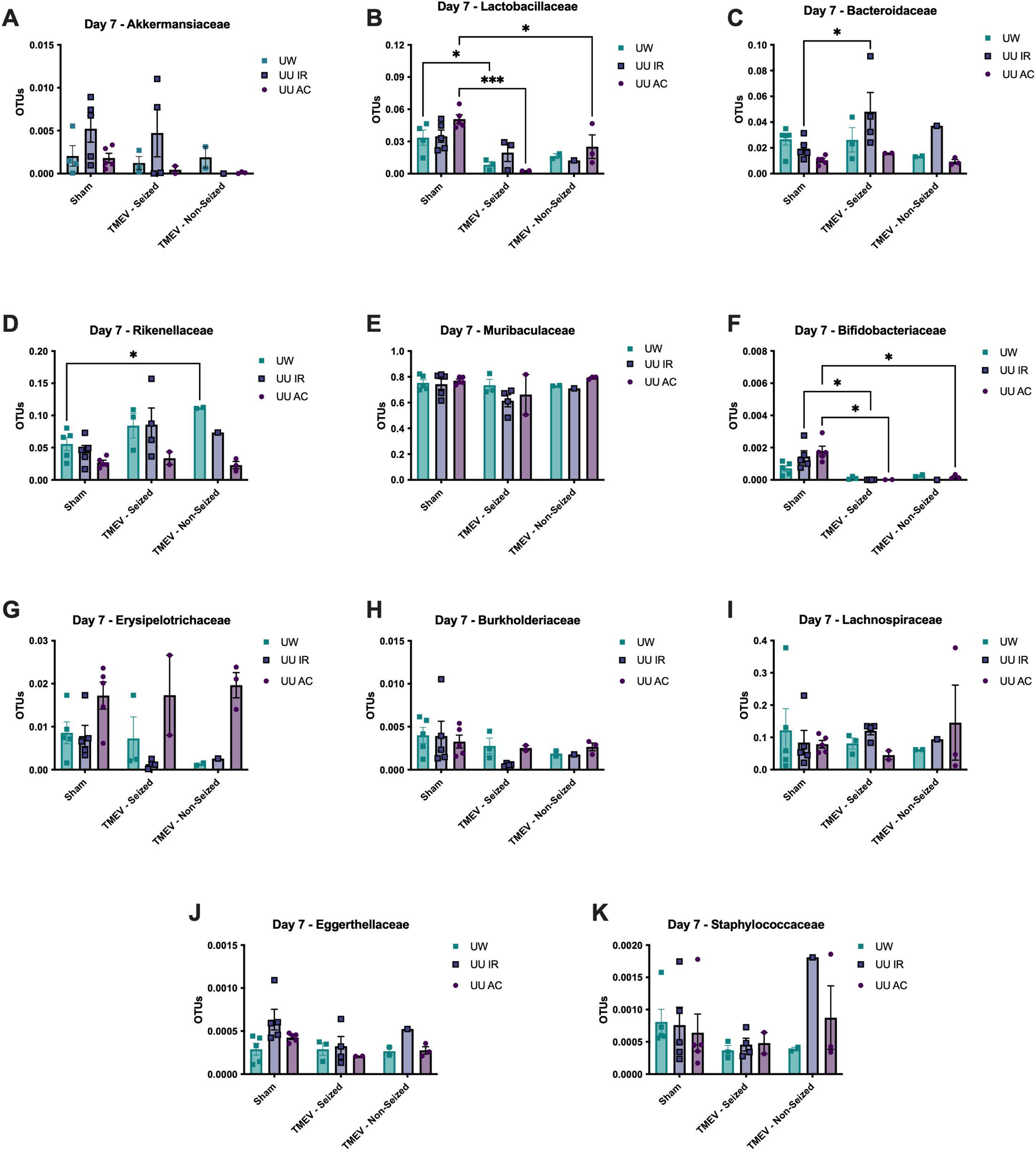
The relative abundance of intestinal microbiome commensal species is impacted by seizure history and dietary condition 7 days after brain infection with the Daniel’s strain of the Theiler’s murine encephalomyelitis virus (TMEV) model in male C57BL/6J mice. There were marked changes in relative abundance of various bacterial species as a function of diet, with the most significant effects observed in Gram-positive bacterial species B) *Lactobacillaceae*, F) *Bifidobacteriaceae*, and J) *Eggerthellaceae* * indicates significant difference, p < 0.05.

### TMEV-infection leads to worsened chronic working memory deficits only in mice fed an autoclaved diet

Acute TMEV infection is known to promote the development of SRS and behavioral comorbidities weeks to months later,^14; 17; 35; 53^ supporting the validity of this approach as a model of infection-induced acquired epilepsy. To determine the extent to which dietary modulation affected onset and severity of behavioral comorbidities of epilepsy, we allowed a subset of mice to recover to 6 weeks p.i. and then monitored for changes in working memory using the novel object place recognition task (NOPR; Figure 6A). Performance in the NOPR task weeks after a TMEV infection can be substantially impaired depending on acute seizure history,^42; 43^ thus we only evaluated the performance of mice infected with TMEV that had at least one handling-induced seizure during the acute infection period. Consistent with our earlier studies, there was a significant main effect of TMEV infection on NOPR performance (F(1,64) = 19.29, p< 0.0001). However, only mice fed the UU AC diet showed significant impairments on NOPR performance relative to diet-matched sham-infected mice (p = 0.0027). Thus, despite seemingly similar acute disease burden during the active TMEV infection period (Figure 2), an autoclaved diet promotes more severe behavioral comorbidities weeks after the acute infection period.

**Figure 6.**
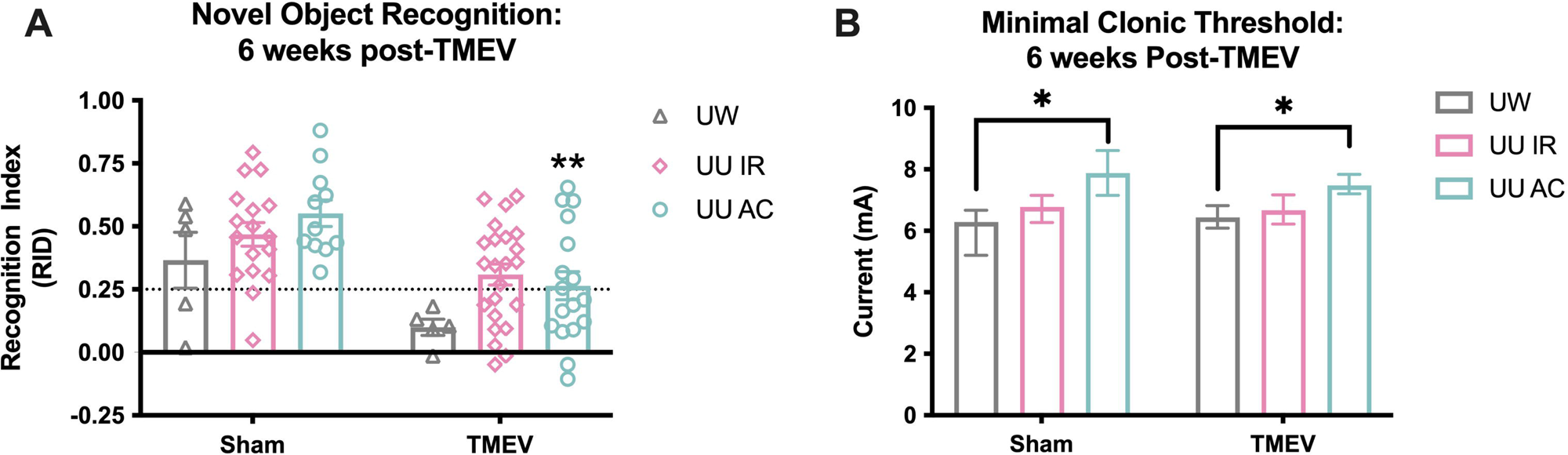
The chronic behavioral changes in male C57B:/6J mice centrally infected with the Daniel’s strain of the Theiler’s murine encephalomyelitis virus (TMEV) was examined 6 weeks post infection. A) While there was a main effect of TMEV infection on NOPR performance (F(1,64) = 19.29, p < 0.0001), the working memory performance was only significantly impaired in TMEV-infected mice fed the autoclaved (AC) UU diet when assessed 6 weeks post-TMEV infection. Neither the UW irradiated (IR) nor UU IR diet-fed mice demonstrated marked cognitive deficits 6 weeks after TMEV infection. ** indicates significantly different from sham-infected, diet-matched mice (p < 0.01). B) The minimal clonic seizure threshold of mice is not itself significantly altered by TMEV infection. However, chronic seizure threshold itself can be substantially influenced by dietary sterilization procedure, as UU AC diet-fed mice demonstrated significantly elevated minimal clonic seizure threshold relative to UW IR diet-fed mice, regardless of TMEV infection status. # Indicates non-overlapping 95% confidence intervals.

### Minimal clonic seizure threshold is markedly altered by dietary formulation but not TMEV infection history

We additionally sought to assess whether diet sterilization influenced the acute seizure threshold of sham- and TMEV-infected mice well after the acute infection period (Figure 6B). The minimal clonic seizure threshold test is a well-characterized model of acute seizure susceptibility that can reveal changes due to pharmacological intervention, genetic manipulation, or neurological insult.^40; 48; 49^ We earlier demonstrated that threshold to an evoked generalized seizure in the i.v. PTZ assay can be substantially altered by acute TMEV infection history, but that this seizure threshold test can evoke significant mortality in C57BL/6J mice.^17^ C57BL/6J mice are generally more susceptible to mortality following a generalized tonic clonic seizure,^54^ thus we sought to use a less severe seizure model to pair to the chronic behavioral testing at 6 weeks after TMEV infection. The minimal clonic test is also known to reveal reduced seizure susceptibility following a TMEV infection.^14^ There were no significant differences between sham- and TMEV-infected mice in the minimal clonic seizure threshold within each dietary group. The UW IR diet-fed mice that were sham infected had a seizure threshold of 6.3 mA [95% confidence intervals - 5.2-6.7] and mice infected with TMEV on the UW IR diet had a seizure threshold of 6.4 mA [6.1-6.8]. The sham-infected UU IR diet-fed mice had a seizure threshold of 6.8 mA [6.3-7.1] and mice infected with TMEV on the UU IR diet had a seizure threshold of 6.7 mA [6.2-7.2]. Sham-infected mice fed the UU AC diet had a seizure threshold of 7.9 mA [7.1-8.6] and mice infected with TMEV on the UU AC diet had a seizure threshold of 7.5 mA [7.2-7.8]. However, there were significant differences in minimal clonic seizure threshold of mice fed the UW IR diet versus the UU AC diet, regardless of TMEV infection status, indicating that dietary sterilization and composition alone can markedly influence chronic seizure threshold and contribute further to potential lab-to-lab variability. However, TMEV infection was not associated with reductions in minimal clonic seizure threshold at 6 weeks post-infection.

## Discussion

This present study demonstrates that differential acute seizure onset and chronic disease burden after TMEV infection can be dramatically altered by changes in dietary sterilization and formulation. The presently observed diet-induced differences in disease phenotype led to substantial changes in the composition of the intestinal microbiome during the acute seizure period. To determine whether the TMEV model may also be sensitive to any diet-induced changes in the composition and diversity of gut commensal microflora, we quantified the changes in the intestinal microbiome prior to and during central infection with TMEV in mice on either matched diet sterilized by autoclaving (AC) or irradiation (IR) versus a different dietary formulation that was IR sterilized (Figure 2). The rationale for these dietary variations was a function of differences intrinsic to two different institutional housing conditions wherein this model was originally described (University of Utah; UU) versus where the subsequent new behavioral studies were performed (University of Washington; UW). Over the course of several years of investigation with numerous cohorts of mice, we presently demonstrate that dietary formulation and sterilization not only significantly shifts the acute handling-induced seizure presentation following central TMEV infection, but also leads to marked differences in intestinal microbiome composition and diversity, as well as stark differences in chronic behavioral comorbidity severity in mice with at least one handling-induced behavioral seizure during the acute viral infection period. Using 16S ribosomal RNA amplicon sequencing of fecal samples collected prior to, during, and post-central infection with TMEV, we herein demonstrate significant shifts from baseline housing (Day −3) in the presence of several bacterial species, including most notably the Gram-positive *Lactobacillaceae, Bifidobacteriaceae,* and *Bacteroides* (Figure 4 and 5). Altogether, the present study points to significant potential for lab-to-lab variability in the disease phenotype of the TMEV mouse model of epilepsy. Our current study reveals a phenomenally unique preclinical model of acquired epilepsy that offers untapped opportunity to critically interrogate the contributions of the gut-brain axis and peripheral immune system on epileptogenesis. Further, our study offers a cautionary note that before embarking on studies using brain infection with TMEV in C567BL/6J mice, it is important to recognize the variability, as discussed below, that can be introduced by changes in extrinsic factors such as diet composition and sterility.

Changes in intestinal commensals can significantly shift anticonvulsant effect in the acute 6 Hz seizure model and *Kcn1a^−/−^*mouse model of genetic epilepsy ^55^, thus our present study also clearly demonstrates that the TMEV model may also be sensitive to depletion of specific intestinal bacteria. Recent studies have also highlighted that key microbial taxa are associated with seizures in the acute infection period of the TMEV model ^56^, albeit that study was limited to evaluations of microbial composition at a single time point 7 DPI from mice housed in different locations. Our studies demonstrate that acute seizures and chronic behavioral biomarkers of chronic disease in the TMEV model are heavily dependent on intestinal microbiome biodiversity. Diet sterilization in TMEV-infected mice residing in a SPF vivarium affects the onset and severity of acute seizures, as well as leads to marked changes in chronic behavioral comorbidities of epilepsy. Further, this study demonstrates that the composition and diversity of the gut microbiome during the active TMEV infection period is markedly altered by diet sterilization procedure, suggesting that the acute and chronic disease phenotype of the TMEV model can be substantially influenced by the intestinal microbiome. This study confirms earlier findings that the acute infection period of the TMEV model is acutely sensitive to dietary modulation,^30^ and extends those studies to highlight that the long-term impact of such dietary manipulation markedly changes the disease trajectory of this mouse model of acquired epilepsy. Our study further reveals that this sensitivity arises at the level of the composition and diversity of the intestinal microbiome (Figures 3, 4, and 5). Importantly, this variability in acute and chronic model phenotype as a result dietary manipulation carries significant implications for the reproducible use of this mouse model for drug discovery applications and points to an important novel therapeutic target in epileptogenesis: namely the gut-brain axis. It has been postulated that the TMEV model of infection-induced epilepsy may identify novel, and as yet, untested therapeutic targets to radically transform the clinical management of epilepsy.^43; 53^ This current study clearly supports the TMEV model as a uniquely differentiated model that is highly variable but altogether provides an unparalleled opportunity to uncover novel molecular targets of relevance to epileptogenesis. Changes in dietary sterilization alone can alter the composition and diversity of enteric microbiota, significantly shifting in the trajectory of both the acute disease burden and chronic epilepsy-related behavioral comorbidities. Our study carries high translational value to advance dietary modification strategies in antiepileptogenesis approaches.

There is reported lab-to-lab variability in the use of the TMEV model. For example, specific viral sub-strains and mouse strains all markedly impact the acute and chronic course of disease presentation in this model.^57^ Indeed, Libbey and colleagues were the first to show that dietary sterilization differences using two unmatched diet formulations could meaningfully alter the acute disease presentation in mice infected with TMEV.^30^ However, that study failed to clearly resolve whether dietary formulation or sterilization were more likely contributing to the observed differences in behavioral seizure presentation during the acute infection period. Our present study aligns with these earlier findings to demonstrate that diet formulation and sterilization differences can markedly change acute disease presentation, but also extends those studies to demonstrate reduced incidence of acute seizures in mice fed an autoclaved versus and a formulation-matched irradiated diet (Figure 2B). Despite this change, there was similar disease burden during the TMEV infection period on days 0-7 p.i. There were also marked differences in body weight gain between AC and IR diet-fed animals (Figure 2A), providing a clear discrepancy in the model reproducibility. Our present work aligns with the findings of Libbey and colleagues to demonstrate that dietary modification can substantially change the acute infection period of the TMEV model. Further, our study supports mounting evidence to point to a potential anticonvulsant and/or disease-modifying role of discrete intestinal commensal species and/or priming of the innate immune system by environmental exposure to modulate epilepsy disease severity.^55^

The variability in this model also highlights an important caveat in the use of the TMEV model for future drug discovery,^53^ and warrants caution in the interpretation of any findings relevant to the discovery of new pharmacological agents. For example, we have previously demonstrated that once-daily oral administration of the ASM, levetiracetam (50 mg/kg), is proconvulsant in TMEV-infected mice fed an irradiated diet, leading to a 50% increase in cumulative seizure burden^53^. However, Metcalf and colleagues subsequently used a nearly identical drug screening protocol to demonstrate that levetiracetam (30 mg/kg, i.p.) is anticonvulsant when delivered from days 3-7 post-infection.^58^ While the diet sterilization procedure was not specifically disclosed in the Metcalf study,^58^ these conflicting reports of the anticonvulsant profile of levetiracetam in the TMEV model indicate that the diversity and composition of the intestinal microbiome alone may vastly influence perceived investigational drug activity in the TMEV model, confounding interpretation of anticonvulsant effects. Indeed, the ASM valproic acid also loses anticonvulsant efficacy against pentylenetetrazol-induced acute seizures with intestinal inflammation modeling ulcerative colitis ^59^, suggesting that the gut microbiome may itself modify ASM activity. Future studies are thus essential to determine the extent to which gut microbiome depletion or modulation alters ASM activity in the TMEV model.

The precise peripheral and central mechanisms underlying our presently observed disease-modifying effects of dietary manipulation in the TMEV model require further scrutiny. It is likely that dietary modulation of the composition and diversity of the intestinal microbiome grossly influences innate immune system response, including the response of resident microglia and infiltrating macrophages. There is less clear evidence that resident CNS microglia cause the acute, handling-induced seizures associated with TMEV infection.^18^ DePaula-Silva and colleagues demonstrated that macrophages infiltrate as early as 3 DPI when acute seizures begin, and peak at 7 days post-infection (DPI), when acute seizure cease.^60^ Further, macrophage infiltration was more substantial in mice with acute seizures than in TMEV-infected mice that did not have acute seizures. Depletion of macrophages can reduce the number of mice with acute seizures^60^; adoptive transfer of monocytes can increase number of mice with seizures. These findings suggest that manipulation of infiltrating macrophages is directly associated with the number of animals with seizures in the TMEV model. DePaula-Silva and colleagues demonstrated that clodronate-mediated depletion of macrophages leads to fewer numbers of mice with seizures following TMEV infection, aligning with findings of Waltl et al 2018.^12; 60^

One impactful clinical strategy to control DRE is through the non-pharmacological ketogenic diet ^61; 62^. However, the mechanisms by which the ketogenic diet specifically mediates any anticonvulsant effect is still intensely investigated ^61–64^. One hypothesized mechanism by which the ketogenic diet can attenuate the burden of epilepsy is through modulation of the diversity and composition of the intestinal microbiome ^25; 55^. As we presently demonstrate, the intestinal microbiome is heavily influenced by dietary intake. Patients with epilepsy can experience significant shifts in the composition and diversity of intestinal microorganisms with adherence to the ketogenic diet ^25^. Preclinical evidence suggests that modification of gut biodiversity by administration of a “ketogenic-like” high fat/low protein diet to wild-type mice housed in a conventional SPF vivarium can significantly elevate acute seizure threshold ^55^. Further, depletion of the intestinal microbiome with antibiotics significantly increased SRS burden in the *Kcna1^−/−^* mouse model of genetic epilepsy ^55^. Administration of selected bacterial commensal species also significantly attenuated SRS in antibiotic-treated *Kcna1^−/−^* mice ^55^. While this particular study was limited to just the use of the acute 6 Hz model of seizure threshold in wild-type mice and the *Kcn1a^−/−^* mouse model of genetic epilepsy ^55^, there is a clear evidence that the intestinal microbiome is an untapped therapeutic modality to modify the severity and burden of epilepsy.

Whether dietary modification or changes in the intestinal microbiome can synergistically improve acute seizure control with available ASMs and/or modify epileptogenesis in acquired, adult-onset epilepsy is clearly in need of further study. The heterogeneity of epilepsy and its related comorbidities, along with the vast number of interactions and variables intrinsic to the intestinal microbiota, make it challenging to conduct the definitive studies to establish the causal relationship between environmental factors and disease course. Our present study demonstrates that modulation of the intestinal microbiome through dietary sterilization may play a role on acute disease presentation after TMEV infection in C57BL/6J male mice and also dramatically alter the trajectory of ensuing disease biomarkers, suggesting that epileptogenesis after TMEV infection in C57BL/6J mice may be altered by dietary therapy or selected bacterial transfer. Remodeling the gut microbiome with probiotics, prebiotics, fecal matter transplantation, ketogenic diet has gained traction in recent years as an untapped opportunity to prevent or treat epilepsy^65–67^. Indeed, short-course antibiotic treatment with the antibiotic ciprofloxacin in adults with drug resistant epilepsy may robustly reduce seizure frequency,^27^ with the *Bacteroidetes: Firmicutes* ratio being substantially altered after treatment. Thus, the potential for synergistic and antagonistic effects of microbiome changes on the activity and efficacy of traditional ASMs in drug-sensitive and drug-resistant individuals, and relevant animal models, is thus in need of further rigorous investigation.

## Author contributions

DKZ, SMM, AE, MBH - conception or design of the work; DKZ, JM, SD, AE, MBH - acquisition, analysis, or interpretation of data; DKZ, HSW, SMM, AE, MBH - have drafted the work or substantively revised it.

## Data Availability

All 16S rRNA amplicon sequencing data supporting the current study are available at the National Center for Biotechnology Information (NCBI) Sequence Read Archive (SRA) as BioProject ID PRJNA1018481.

## Literature Cited

1. Singh G, Prabhakar S. The association between central nervous system (CNS) infections and epilepsy: epidemiological approaches and microbiological and epileptological perspectives. Epilepsia 2008;49 Suppl 6:2–7.

2. Khetsuriani N, Holman RC, Anderson LJ. Burden of encephalitis-associated hospitalizations in the United States, 1988-1997. Clin Infect Dis 2002;35:175–182.

3. Ngarka L, Siewe Fodjo JN, Aly E, Masocha W, Njamnshi AK. The Interplay Between Neuroinfections, the Immune System and Neurological Disorders: A Focus on Africa. Front Immunol 2021;12:803475.

4. Carpio A, Hauser W, Aguirre R, Roman M. Etiology of epilepsy in Ecuador. Epilepsia 2001;42.

5. Annegers JF, Hauser WA, Lee JR, Rocca WA. Incidence of acute symptomatic seizures in Rochester, Minnesota, 1935-1984. Epilepsia 1995;36:327–333.

6. Ba-Diop A, Marin B, Druet-Cabanac M, Ngoungou EB, Newton CR, Preux PM. Epidemiology, causes, and treatment of epilepsy in sub-Saharan Africa. Lancet Neurol 2014;13:1029–1044.

7. Antony AR, Haneef Z. Systematic review of EEG findings in 617 patients diagnosed with COVID-19. Seizure 2020;83:234–241.

8. Khedr EM, Shoyb A, Mohammaden M, Saber M. Acute symptomatic seizures and COVID-19: Hospital-based study. Epilepsy Res 2021;174:106650.

9. Misra UK, Tan CT, Kalita J. Viral encephalitis and epilepsy. Epilepsia 2008;49 Suppl 6:13–18.

10. Annegers JF, Hauser WA, Beghi E, Nicolosi A, Kurland LT. The risk of unprovoked seizures after encephalitis and meningitis. Neurology 1988;38:1407–1410.

11. Cruzado D, Masserey-Spicher V, Roux L, Delavelle J, Picard F, Haenggeli CA. Early onset and rapidly progressive subacute sclerosing panencephalitis after congenital measles infection. Eur J Pediatr 2002;161:438–441.

12. Waltl I, Kaufer C, Broer S, Chhatbar C, Ghita L, Gerhauser I, et al. Macrophage depletion by liposome-encapsulated clodronate suppresses seizures but not hippocampal damage after acute viral encephalitis. Neurobiol Dis 2018;110:192–205.

13. Waltl I, Kaufer C, Gerhauser I, Chhatbar C, Ghita L, Kalinke U, et al. Microglia have a protective role in viral encephalitis-induced seizure development and hippocampal damage. Brain Behav Immun 2018;74:186–204.

14. Stewart KA, Wilcox KS, Fujinami RS, White HS. Theiler’s virus infection chronically alters seizure susceptibility. Epilepsia 2010;51:1418–1428.

15. Barker-Haliski ML, Dahle EJ, Heck TD, Pruess TH, Vanegas F, Wilcox KS, et al. Evaluating an Etiologically-Relevant Platform for Therapy Development for Temporal Lobe Epilepsy: Effects of Carbamazepine and Valproic Acid on Acute Seizures and Chronic Behavioral Comorbidities in the Theiler’s Murine Encephalomyelitis Virus Mouse Model. J Pharmacol Exp Ther 2015.

16. Kaufer C, Chhatbar C, Broer S, Waltl I, Ghita L, Gerhauser I, et al. Chemokine receptors CCR2 and CX3CR1 regulate viral encephalitis-induced hippocampal damage but not seizures. Proc Natl Acad Sci U S A 2018;115:E8929–E8938.

17. Barker-Haliski ML, Heck TD, Dahle EJ, Vanegas F, Pruess TH, Wilcox KS, et al. Acute treatment with minocycline, but not valproic acid, improves long-term behavioral outcomes in the Theiler’s virus model of temporal lobe epilepsy. Epilepsia 2016;57:1958–1967.

18. Cusick MF, Libbey JE, Patel DC, Doty DJ, Fujinami RS. Infiltrating macrophages are key to the development of seizures following virus infection. J Virol 2013;87:1849–1860.

19. Patel DC, Wallis G, Dahle EJ, McElroy PB, Thomson KE, Tesi RJ, et al. Hippocampal TNFalpha Signaling Contributes to Seizure Generation in an Infection-Induced Mouse Model of Limbic Epilepsy. eNeuro 2017;4.

20. Libbey JE, Kennett NJ, Wilcox KS, White HS, Fujinami RS. Interleukin-6, produced by resident cells of the central nervous system and infiltrating cells, contributes to the development of seizures following viral infection. J Virol 2011;85:6913–6922.

21. Auvin S, Shin D, Mazarati A, Sankar R. Inflammation induced by LPS enhances epileptogenesis in immature rat and may be partially reversed by IL1RA. Epilepsia 2010;51 Suppl 3:34–38.

22. Auvin S, Mazarati A, Shin D, Sankar R. Inflammation enhances epileptogenesis in the developing rat brain. Neurobiol Dis 2010;40:303–310.

23. Mazarati A, Shin D, Auvin S, Caplan R, Sankar R. Kindling epileptogenesis in immature rats leads to persistent depressive behavior. Epilepsy Behav 2007;10:377–383.

24. Mendes V, Galvao I, Vieira AT. Mechanisms by Which the Gut Microbiota Influences Cytokine Production and Modulates Host Inflammatory Responses. J Interferon Cytokine Res 2019;39:393–409.

25. Hampton T. Gut Microbes May Account for the Anti-Seizure Effects of the Ketogenic Diet. JAMA 2018;320:1307.

26. Gomez-Eguilaz M, Ramon-Trapero JL, Perez-Martinez L, Blanco JR. The beneficial effect of probiotics as a supplementary treatment in drug-resistant epilepsy: a pilot study. Benef Microbes 2018;9:875–881.

27. Cheraghmakani H, Rezai MS, Valadan R, Rahimzadeh G, Moradi M, Jahanfekr V, et al. Ciprofloxacin for treatment of drug-resistant epilepsy. Epilepsy Res 2021;176:106742.

28. Gao X, Cao Q, Cheng Y, Zhao D, Wang Z, Yang H, et al. Chronic stress promotes colitis by disturbing the gut microbiota and triggering immune system response. Proc Natl Acad Sci U S A 2018;115:E2960–E2969.

29. Raible DJ, Frey LC, Del Angel YC, Carlsen J, Hund D, Russek SJ, et al. JAK/STAT pathway regulation of GABAA receptor expression after differing severities of experimental TBI. Exp Neurol 2015;271:445–456.

30. Libbey JE, Doty DJ, Sim JT, Cusick MF, Round JL, Fujinami RS. The effects of diet on the severity of central nervous system disease: One part of lab-to-lab variability. Nutrition 2016;32:877–883.

31. Meeker S, Beckman M, Knox KM, Treuting PM, Barker-Haliski M. Repeated Intraperitoneal Administration of Low-Concentration Methylcellulose Leads to Systemic Histologic Lesions Without Loss of Preclinical Phenotype. J Pharmacol Exp Ther 2019.

32. Kilkenny C, Browne W, Cuthill IC, Emerson M, Altman DG, National Centre for the Replacement R, et al. Animal research: reporting in vivo experiments--the ARRIVE guidelines. J Cereb Blood Flow Metab 2011;31:991–993.

33. Barker-Haliski ML, Dahle EJ, Heck TD, Pruess TH, Vanegas F, Wilcox KS, et al. Evaluating an etiologically relevant platform for therapy development for temporal lobe epilepsy: effects of carbamazepine and valproic acid on acute seizures and chronic behavioral comorbidities in the Theiler’s murine encephalomyelitis virus mouse model. J Pharmacol Exp Ther 2015;353:318–329.

34. Libbey JE, Kirkman NJ, Smith MC, Tanaka T, Wilcox KS, White HS, et al. Seizures following picornavirus infection. Epilepsia 2008;49:1066–1074.

35. Stewart KA, Wilcox KS, Fujinami RS, White HS. Development of postinfection epilepsy after Theiler’s virus infection of C57BL/6 mice. J Neuropathol Exp Neurol 2010;69:1210–1219.

36. Racine RJ. Modification of seizure activity by electrical stimulation: II. Motor seizure. Electroenceph. Clin. Neurophysiol. 1972;32:281–294.

37. Loewen JL, Barker-Haliski ML, Dahle EJ, White HS, Wilcox KS. Neuronal Injury, Gliosis, and Glial Proliferation in Two Models of Temporal Lobe Epilepsy. J Neuropathol Exp Neurol 2016;75:366–378.

38. Dunham MS, Miya TA. A note on a simple apparatus for detecting neurological deficit in rats and mice. J. Amer. Pharm. Ass. Sci. Ed. 1957;46:208–209.

39. Ericsson AC, Davis JW, Spollen W, Bivens N, Givan S, Hagan CE, et al. Effects of vendor and genetic background on the composition of the fecal microbiota of inbred mice. PLoS One 2015;10:e0116704.

40. Beckman M, Knox K, Koneval Z, Smith C, Jayadev S, Barker-Haliski M. Loss of presenilin 2 age-dependently alters susceptibility to acute seizures and kindling acquisition. Neurobiol Dis 2020;136:104719.

41. Barker-Haliski ML, Vanegas F, Mau MJ, Underwood TK, White HS. Acute cognitive impact of antiseizure drugs in naive rodents and corneal-kindled mice. Epilepsia 2016;57:1386–1397.

42. Umpierre AD, Remigio GJ, Dahle EJ, Bradford K, Alex AB, Smith MD, et al. Impaired cognitive ability and anxiety-like behavior following acute seizures in the Theiler’s virus model of temporal lobe epilepsy. Neurobiol Dis 2014;64:98–106.

43. Barker-Haliski M, Nishi T, White HS. Soticlestat, a novel cholesterol 24-hydroxylase inhibitor, modifies acute seizure burden and chronic epilepsy-related behavioral deficits following Theiler’s virus infection in mice. Neuropharmacology 2023;222:109310.

44. Hammond RS, Tull LE, Stackman RW. On the delay-dependent involvement of the hippocampus in object recognition memory. Neurobiol Learn Mem 2004;82:26–34.

45. Brandt C, Tollner K, Klee R, Broer S, Loscher W. Effective termination of status epilepticus by rational polypharmacy in the lithium-pilocarpine model in rats: Window of opportunity to prevent epilepsy and prediction of epilepsy by biomarkers. Neurobiol Dis 2015;75:78–90.

46. Barker-Haliski M, Harte-Hargrove LC, Ravizza T, Smolders I, XIao B, Brandt C, et al. A Companion to the Preclinical Common Data Elements for Pharmacological Studies in Animal Models of Seizures and Epilepsy. A Report of the TASK3 Pharmacology Working Group of the ILAE/AES Joint Translational Task Force. Epilepsia Open 2018;In press.

47. Beckman M, Knox K, Koneval Z, Smith C, Jayadev S, Barker-Haliski M. Loss of presenilin 2 age-dependently alters susceptibility to acute seizures and kindling acquisition. Neurobiol Dis 2019;136:104719.

48. Otto JF, Singh NA, Dahle EJ, Leppert MF, Pappas CM, Pruess TH, et al. Electroconvulsive seizure thresholds and kindling acquisition rates are altered in mouse models of human KCNQ2 and KCNQ3 mutations for benign familial neonatal convulsions. Epilepsia 2009;50:1752–1759.

49. Singh NA, Otto JF, Dahle EJ, Pappas C, Leslie JD, Vilaythong A, et al. Mouse models of human KCNQ2 and KCNQ3 mutations for benign familial neonatal convulsions show seizures and neuronal plasticity without synaptic reorganization. J Physiol 2008;586:3405–3423.

50. Bliss CI. The method of probits. Science 1934;79:38–39.

51. Cryan JF, O’Riordan KJ, Cowan CSM, Sandhu KV, Bastiaanssen TFS, Boehme M, et al. The Microbiota-Gut-Brain Axis. Physiol Rev 2019;99:1877–2013.

52. Pickard JM, Zeng MY, Caruso R, Nunez G. Gut microbiota: Role in pathogen colonization, immune responses, and inflammatory disease. Immunol Rev 2017;279:70–89.

53. Barker-Haliski ML, Loscher W, White HS, Galanopoulou AS. Neuroinflammation in epileptogenesis: Insights and translational perspectives from new models of epilepsy. Epilepsia 2017;58 Suppl 3:39–47.

54. Collard R, Aziz MC, Rapp K, Cutshall C, Duyvesteyn E, Metcalf CS. Galanin analogs prevent mortality from seizure-induced respiratory arrest in mice. Front Neural Circuits 2022;16:901334.

55. Olson CA, Vuong HE, Yano JM, Liang QY, Nusbaum DJ, Hsiao EY. The Gut Microbiota Mediates the Anti-Seizure Effects of the Ketogenic Diet. Cell 2018;174:497.

56. Gallucci A, Patel DC, Thai K, Trinh J, Gude R, Shukla D, et al. Gut metabolite S-equol ameliorates hyperexcitability in entorhinal cortex neurons following Theiler murine encephalomyelitis virus-induced acute seizures. Epilepsia 2021;62:1829–1841.

57. Broer S, Kaufer C, Haist V, Li L, Gerhauser I, Anjum M, et al. Brain inflammation, neurodegeneration and seizure development following picornavirus infection markedly differ among virus and mouse strains and substrains. Exp Neurol 2016;279:57–74.

58. Metcalf CS, Vanegas F, Underwood T, Johnson K, West PJ, Smith MD, et al. Screening of prototype antiseizure and anti-inflammatory compounds in the Theiler’s murine encephalomyelitis virus model of epilepsy. Epilepsia Open 2022;7:46–58.

59. De Caro C, Leo A, Nesci V, Ghelardini C, di Cesare Mannelli L, Striano P, et al. Intestinal inflammation increases convulsant activity and reduces antiepileptic drug efficacy in a mouse model of epilepsy. Sci Rep 2019;9:13983.

60. DePaula-Silva AB, Sonderegger FL, Libbey JE, Doty DJ, Fujinami RS. The immune response to picornavirus infection and the effect of immune manipulation on acute seizures. J Neurovirol 2018;24:464–477.

61. Rho JM. How does the ketogenic diet induce anti-seizure effects? Neurosci Lett 2017;637:4–10.

62. Simeone TA, Simeone KA, Rho JM. Ketone Bodies as Anti-Seizure Agents. Neurochem Res 2017.

63. Clanton RM, Wu G, Akabani G, Aramayo R. Control of seizures by ketogenic diet-induced modulation of metabolic pathways. Amino Acids 2017;49:1–20.

64. Chang P, Augustin K, Boddum K, Williams S, Sun M, Terschak JA, et al. Seizure control by decanoic acid through direct AMPA receptor inhibition. Brain 2016;139:431–443.

65. Iannone LF, Gomez-Eguilaz M, De Caro C. Gut microbiota manipulation as an epilepsy treatment. Neurobiol Dis 2022;174:105897.

66. Mejia-Granados DM, Villasana-Salazar B, Lozano-Garcia L, Cavalheiro EA, Striano P. Gut-microbiota-directed strategies to treat epilepsy: clinical and experimental evidence. Seizure 2021;90:80–92.

67. Kundu S, Nayak S, Rakshit D, Singh T, Shukla R, Khatri DK, et al. The microbiome-gut-brain axis in epilepsy: pharmacotherapeutic target from bench evidence for potential bedside applications. Eur J Neurol 2023.

